# A Conserved Requirement for RME-8/DNAJC13 in Neuronal Autolysosome Reformation

**DOI:** 10.1101/2023.02.27.530319

**Authors:** Sierra Swords, Nuo Jia, Anne Norris, Jil Modi, Qian Cai, Barth D. Grant

## Abstract

Autophagosomes fuse with lysosomes, forming autolysosomes that degrade engulfed cargo. To maintain lysosomal capacity, autolysosome reformation (ALR) must regenerate lysosomes from autolysosomes using a membrane tubule-based process. Maintaining lysosomal capacity is required to maintain proteostasis and cellular health, especially in neurons where lysosomal dysfunction has been repeatedly implicated in neurodegenerative disease. Cell biological studies have linked the DNA-J domain Hsc70 co-chaperone RME-8/DNAJC13 to endosomal coat protein regulation, while human genetics studies have linked RME-8/DNAJC13 to neurological disease, including Parkinsonism and Essential Tremor. We report new analysis of the requirements for the RME-8/DNAJC13 protein in neurons, focusing on *C. elegans* mechanosensory neurons in the intact animal, and in primary mouse cortical neurons in culture. We find that loss of RME-8/DNAJC13 in both systems results in accumulation of grossly elongated autolysosomal tubules. Further *C. elegans* analysis revealed a similar autolysosome tubule accumulation defect in mutants known to be required for ALR in mammals, including bec-1/beclin and vps-15/PIK3R4/p150 that regulate type-III PI3-kinase VPS-34, and dyn-1*/dynamin* that severs ALR tubules. Clathrin is also an important ALR regulator implicated in autolysosome tubule formation and release. In *C. elegans* we found that loss of RME-8 causes severe depletion of clathrin from neuronal autolysosomes, a phenotype shared with *bec-1* and *vps-15* mutants. We conclude that RME-8/DNAJC13 plays a conserved but previously unrecognized role in autolysosome reformation, likely affecting ALR tubule initiation and/or severing. Additionally, in both systems, we found that loss of RME-8/DNAJC13 appeared to reduce autophagic flux, suggesting feedback regulation from ALR to autophagy. Our results connecting RME-8/DNAJC13 to ALR and autophagy provide a potential mechanism by which RME-8/DNAJC13 could influence neuronal health and the progression of neurodegenerative disease.

## INTRODUCTION

Organelle integrity and cellular proteostasis depend heavily upon a variety of degradative mechanisms for macromolecular turnover. Among these, one key pathway is macroautophagy (hereafter referred to as autophagy), a mechanism that can remove whole defective organelles and cytoplasmic protein aggregates as they build up, ameliorating the cellular dysfunction that can occur if such components accumulate. Autophagy begins with the phagophore, a *de novo* membrane which expands to completely engulf a region of cytoplasm, often including a specific degradative target, forming a double-membraned autophagosome. Once autophagosomes form, they fuse with lysosomes, creating autolysosomes to degrade their contents.

Lysosomes are reformed from autolysosomes using a specific mechanism, termed autolysosome reformation (ALR)[1]. This autolysosome reformation (ALR) process is required to maintain the active lysosome pool, allowing degradation of autophagic contents to continue uninterrupted[1]. During ALR, recycling tubules are drawn out from buds on the autolysosome limiting membrane and are then severed to release protolysosomes. The mechanism of ALR tubule formation and release is not well understood, but appears to require phosphoinositide lipids P1(3)P and P1(4,5)P_2_, a clathrin coat, the severing enzyme dynamin, and kinesin motor autolysosomes bearing grossly elongated membrane tubules. In mouse cortical neurons, we also found that depletion of RME-8/DNAJC13 resulted in gross enlargement of the autolysosomes, and loss of most LC3-negative lysosomes. Further *C. elegans* analysis revealed a similar autolysosome tubule accumulation defect in mutants known to be required for ALR in mammals, including *bec-1/beclin and vps-15/PIK3R4/p150* that regulate the VPS-34 type-III PI3-kinase, and *dyn-1/dynamin* that severs ALR tubules. Furthermore, we found that loss of RME-8 causes severe depletion of clathrin from autolysosomes, a phenotype also shared with *snx-1, bec-1* and *vps-15* mutants. Taken together, we conclude that RME-8/DNAJC13 plays a conserved but previously unrecognized role in autolysosome reformation, likely affecting ALR tubule severing. Additionally, in both systems, we found that loss of RME-8/DNAJC13 reduced autophagic flux, suggesting feedback regulation from ALR to autophagy. These results connecting RME-8/DNAJC13 to ALR and autophagy provide a potential mechanism by which RME-8/DNAJC13 could influence neuronal health and the progression of neurodegenerative diseases, including Parkinson’s Disease.

## RESULTS

### Loss of RME-8 function results in accumulation of long lysosomal tubules

Given the association of RME-8/DNAJC13 with neurological disease[13, 17, 29–32], we sought to better understand RME-8 function in neurons of *C. elegans* where RME-8-mediated trafficking mechanisms were first elucidated[15, 19, 23, 24]). Here we focused on the mechanosensory touch neurons, in particular analyzing the two centrally located touch neurons referred to as ALM right and ALM left (Figure 1A). All six touch neurons are embedded in the hypodermis (skin), very close to the cuticle, where they sense gentle mechanical stimuli[33]. The stereotyped location of the ALM neurons close to the exterior of the animal, and their simple architecture with one main process that extends anteriorly to make synapses in the head, make these neurons ideal for quantitative imaging in the living, intact animal. To allow subcellular analysis specifically in these neurons, we expressed a variety of fluorescently tagged proteins from single copy transgenes, driven by the mechanosensory touch neuron-specific *mec-7* promoter[34].

**Figure 1.**
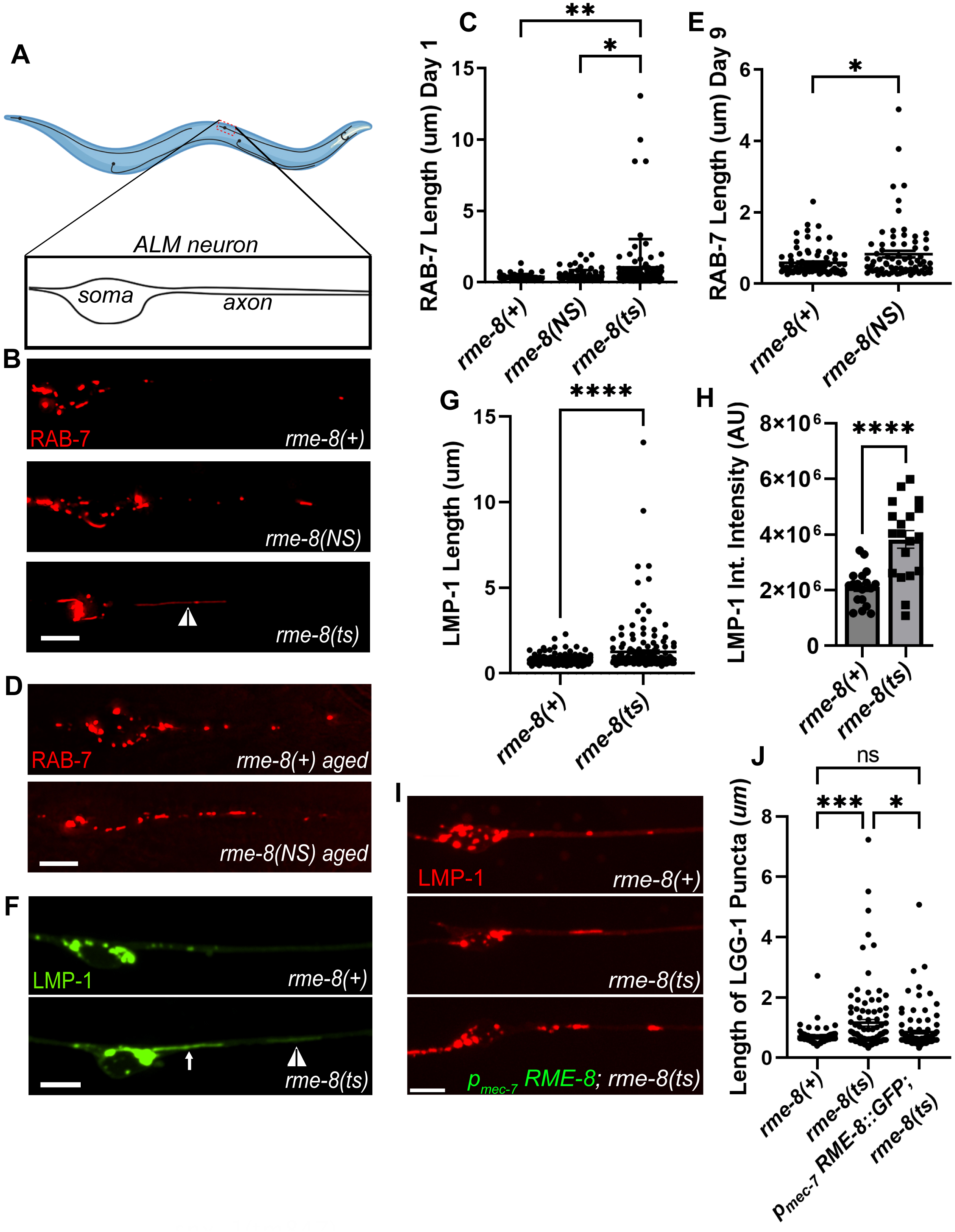
Loss of RME-8 results in accumulation of elongated lysosomal tubules. **A)** Representative drawing of ALM neurons in *C. elegans.* One of two ALM neurons is shown (other is located laterally opposite). Black dashed box represents the approximate area that is shown in example micrographs. **B)** Fluorescent micrographs showing pmec-7 promoter-driven mScarletl::RAB-7 labeled (late endosome/ lysosome marker) puncta in soma (left) and proximal axon (right) in ALM neurons in *rme-8(+), rme-8(N861S),* and *rme-8(b1023ts)* Day 1 adult backgrounds. White arrowhead indicates grossly elongated tubule. **C)** Length of RAB-7 puncta/ tubule in the proximal axon in *rme-8(+), rme-8(N861S)* and *rme-8(b1023ts)* is graphed in Day 1 adult animals. Each data point represents an individual puncta/tubule. A minimum of 15 animals were analyzed per strain. **D)** Pmec-7 promoter-driven mScarletl::RAB-7 labeled puncta in *rme-8(+)* and *rme-8(N861S)* backgrounds in Day 9 adult. **E)** Length of RAB-7 puncta/ tubule in the proximal axon in *rme-8(+),* and *rme-8(N861S)* is graphed in Day 9 adults. Each data point represents an individual puncta/tubule. **F)** Pmec-7 LMP-1::mNeonGreen (NG) labeled (lysosome marker) puncta. A severely elongated LMP-1 positive tubule can be seen emanating into the axon from puncta in the soma (white arrow), and another dimmer tubule can be seen further out in the axon (white arrowhead). **G)** Quantification of LMP-1::NG axonal puncta/ tubule length seen in conditions shown in E). Each data point represents an individual puncta/tubule. A minimum of 15 animals were analyzed per strain. **H)** Graph depicting LMP-1::mNG Integrated intensity (Arbitrary Units; AU) for the same experiment as 1E and 1F. Each data point represents the integrated intensity of all thresholded puncta in one soma in one animal. **I)** Micrographs of pmec-7 RME-8::oxGFP rescue of LMP-1::mScarleti length in *rme-8(b1023ts)* background. **J)** Graph quantifying elongation of LMP-1::mScarletl in *rme-8(+), rme-8(b1023ts),* and *rme-8(b1023ts); pmec-7 RME-8::GFP.* Scale bars, 5 um. Abbreviations: *rme-8(NS)= rme-8(pw22[N8615]); rme-8(ts)= rme-8(b1023ts)*. **C), E),** and **J):** One-way ANOVA followed with Tukey’s Multiple Comparisons Test. **G), H):** Student’s unpaired t-test. *P < 0.05, **P < 0.01, ***P < 0.001, ****P < 0.0001.

*rme-8* is an essential gene, so for most experiments we measured the effects of loss of RME-8 function on neuronal trafficking using the well-characterized temperature sensitive allele, *rme­8(b1023ts)*[19]. Temperature shifting this strain from the permissive temperature (15°C) to the non-permissive temperature (25°C) is known to destabilize the RME-8 protein, and results in clear phenotypes within 24 hours[19]. For most experiments we temperature shifted *rme­8(b1023ts)* animals at the L4 stage, and conducted imaging experiments in young adults 24-30 hours later. For some experiments we also analyzed the effects of *rme-8(pw22[N861S]),* an allele we generated by CRISPR-based genome engineering to mimic the human late onset Parkinson’s associated allele N8555[13]. *rme-8(pw22[N861S])* bears an equivalent point mutation to N855S, changing asparagine 861 to serine (N8615).

Given the known roles of RME-8 on endosomes, and the association of neurodegenerative disease with membrane trafficking pathway dysfunction, we initially analyzed the effects of the *rme-8(ts)* mutant on late endosomes and lysosomes using an mScarlet::RAB-7 marker expressed in the mechanosensory neurons. Importantly, in young adult (adult Day 1) *rme-8(b1023ts)* mutants we noted the appearance of RAB-7 vesicles bearing highly elongated tubules, a morphology not observed in control animals (Figure 1B, white arrowhead; quantified in 1C). We also assayed for this phenotype in the PD-allele mimic *rme-8(N861S).* Interestingly, we observed a similar accumulation of elongated RAB-7-positive tubules in aged animals in late adulthood (adult Day 9), but not in young adults (Figure 1D-E). This mimics the late-onset nature of the homologous PD-linked allele in human patients, suggesting that the N861S allele confers a similar but more subtle defect than the *b1023ts* loss of function allele. Tubules emanating from endosomes and lysosomes are generally associated with recycling processes, with accumulation of exaggerated tubules often associated with defects in tubule release.

To better define the nature of these vesicles, we extended our analysis using LMP-1/LAMP1 fused to mNeonGreen (mNG), a transmembrane marker of lysosomes (Figure 1F). We noted a very similar phenotype using the lysosome marker in *rme-8(b1023ts)* mutants, measuring a highly significant increase in LMP-1 tubule length (Figure 1G). We noted that in many cases these long thin tubules appeared connected to a larger vesicle located in the neuronal soma (white arrow, Figure 1F; Video S1), and in others the vesicle and tubule were located in the soma-proximal neuronal process (white arrowhead, Figure 1F). These persistent tubules were not seen in *rme-8(+)* backgrounds (Figure 1F, Video S2). We extended this analysis, measuring the intensity LMP-1::mNG labeling of lysosomes in the soma, dendrite, and proximal axon. Interestingly, LMP-1 integrated intensity was increased upon loss of RME-8, indicating a significant abnormal accumulation of LMP-1 labeled lysosomes within neurons upon loss of RME-8 (Figure 1H). These data indicate a key requirement for RME-8 in neuronal lysosome homeostasis.

### RME-8 function is touch neuron autonomous

To confirm that the lysosomal tubule phenotype is caused by loss of RME-8, and test if RME-8 is required autonomously within the touch neurons to maintain lysosome homeostasis, we tested for rescue of the neuronal *rme-8(b1023ts)* phenotype in animals expressing a touch neuron-specific *rme-8(+)* minigene. Indeed, we observed that expressing wild-type RME-8 in just the six touch neurons rescued touch neuron lysosomal tubule accumulation (Figure 1I-J).

### RME-8 affects autolysosomes

Lysosomes receive cargo via membrane fusion, fusing with endosomes, phagosomes, and autophagosomes to form endolysosomes, phagolysosomes, and autolysosomes respectively (reviewed in [35–37]). Neurons in particular display high levels of basal autophagy compared to other cell types[8]. Thus, we sought to determine to what extent the LMP-1 positive structures we observed in the touch neurons represent autolysosomes. To achieve this, we created strains expressing the autophagosome marker mNG::LGG-1 and LMP-1::mScarlet-I (mSc) in touch neurons (Figure 2A). We measured a 75% pixel overlap of mNG::LGG-1 and LMP-1::mSc, with virtually all LMP-1-positive lysosomes containing at least some mNG::LGG-1, indicating that they are autolysosomes (Figure 2B). We only rarely observed mNG::LGG-1 labeled objects lacking LMP-1::mSc (Figure 2A, white arrow), indicating that at steady state most autophagosomes have already fused with lysosomes.

**Figure 2.**
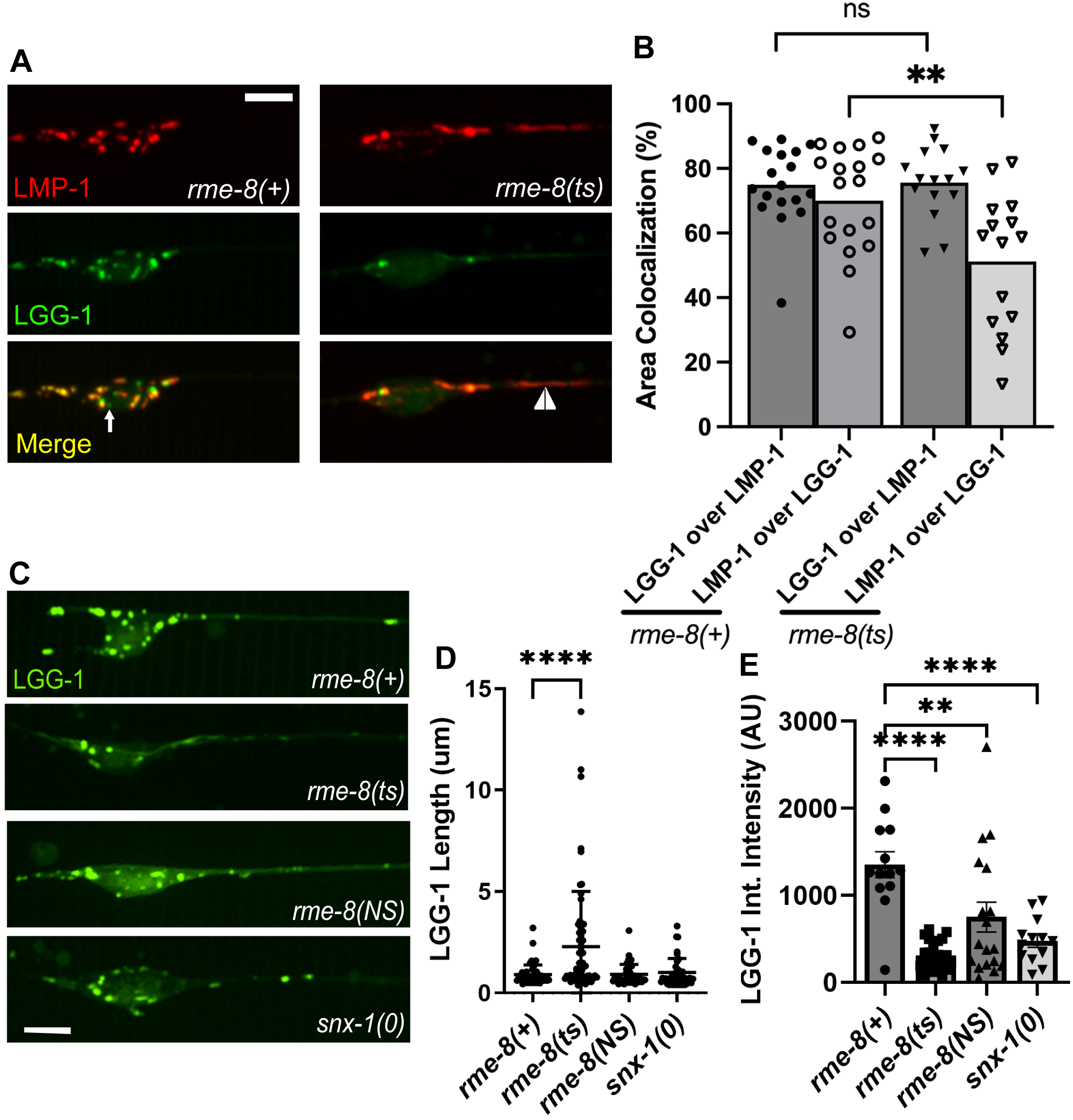
*rme-8* mutants accumulate abnormal autolysosomes. **A)** Micrographs are shown for single channels as well as merged images in double-labeled LMP-1::mScarlet-1(lysosome marker); mNG::LGG-1 (autophagosome marker) in *rme-8(+)* and *rme-8(b1023ts)* backgrounds. White arrow labels mNG::LGG-1+/ LMP-1::mSc- puncta. White arrowhead labels tubules labeled by LMP-1 and dimly labeled by LGG-1 can be seen in the *rme­8(b1023ts)* background. **B)** Graphs quantifying the colocalization of LMP-1:mSc and mNG::LGG-1, measured by %Area Colocalization. Each data point represents average % Area colocalization for one animal. **P < 0.01, ns > 0.05 by two-tailed unpaired t-test. **C)** Micrograph showing pmec-7::mNG::LGG-1 in neurons of *rme-8(+), rme-8(b1023ts), rme-8(N861S),* and *snx-1(tm847)* backgrounds. Scale bar: 5um. **D)** Graph quantifying elongation of LGG-1::mNG in *rme-8(+), rme-8(b1023ts),* and *rme-8(N861S); snx-1(0)* backgrounds. Each data point represents an individual puncta/tubule. ****P < 0.0001 by One-way ANOVA followed with Tukey’s Multiple Comparisons Test. **E)** Graph depicting mNG::LGG-1 Integrated intensity (Arbitrary Units; AU) for the same experiment as 2C and 2D. Each data point represents the integrated intensity of all thresholded puncta in one soma in one animal. ****P < 0.0001, **P<0.01 by One-way ANOVA followed with Tukey’s Multiple Comparisons Test.

We observed the grossly elongated tubules (white arrowhead) in *rme-8(b1023ts)* mutants in LMP-1::mSc; mNG::LGG-1 double labeled strains (Figure 2A) and in single labeled mNG::LGG-1 strains (Figure 2C, quantified in 2D). We noted that the overall mNG::LGG-1 signal was weaker in *rme-8* mutants, with mNG::LGG-1 fluorescence particularly weak and difficult to detect in the tubules. mNG::LGG-1 fluorescence was much more prominent in vesicular region of the neuronal autolysosomes (observed in Figures 2A, 2C). This was reflected in a maintained 76% overlap of LGG-1-positive area that colocalized with LMP-1 in *rme-8(b1023ts)* mutants, but a reduced percentage of LMP-1-positive area that colocalized with LGG-1 signal in *rme­8(b1023ts)* mutants (Figure 2B). We also measured a reduced overall integrated intensity for LGG-1::mNG puncta in the neuronal soma in *rme-8(b1023ts)* mutants (Figure 2E). Taken together our results indicate that *C. elegans* touch neurons maintain active autophagy under well-fed conditions, and most lysosomes in the neuronal soma have previously fused with autophagosomes, with loss of RME-8 leading to accumulation of abnormal autolysosome­derived membrane tubules.

### Loss of RME-8 phenocopies autolysosome reformation mutants

The strikingly elongated LMP-1-positive tubules emanating from autolysosome puncta were highly reminiscent of autolysosome reformation tubules as reported in mammalian cells[1, 2, 4, 6, 38]. ALR tubules will persist and elongate if the mechanism by which the tubules are severed is impaired[2, 4, 6, 38, 39]. Our observations that LMP-1/LAMP1 intensity in the autolysosomes increased in *rme-8* mutants suggests a failure to recycle LMP-1 out of the autolysosome, further supporting the hypothesis that the elongated tubules accumulating in *rme-8* mutant neurons represent an accumulating ALR intermediate (Figure 2). Similar effects have been observed in other organisms when ALR was impaired[38].

If the *rme-8* mutant phenotype indicates a defect in neuronal ALR, we would expect to find similar phenotypes in *C. elegans* mutants lacking proteins previously identified as ALR regulators in other systems. In particular, enzymes that regulate lysosomal phosphatidylinositol 4,5-bisphosphate (PI(4,5)P_2_), phosphatidylinositol 3-phosphate (PI(3)P), as well as the coat protein clathrin and the membrane severing GTPase dynamin, have been suggested to function in ALR, in addition to their more well-known roles in other trafficking steps [2, 3, 6]. For instance, mammalian cells inhibited for class III phosphoinositide 3-kinase VPS34 activity display lysosomal tubule accumulation during ALR [6]. To impair the VPS-34 complex in *C. elegans,* we used mutants in *bec-1/beclin* and *vps-15/PIK3R4/p150,* key regulators of VPS-34 activity[40–42]. We observed severely elongated LMP-1::mNG tubules in *bec-1* and *vps-15* mutants that strongly resemble those we found in *rme-8* mutants (Figure 3A, white arrowheads, quantified in 3B). We also examined the tubule accumulation phenotype in neurons impaired for dynamin[43]. Previous work showed that mammalian dynamin 2 binds to autolysosome tubules along their length, and pharmacologic inhibition of dynamin 2 in hepatocytes produced enlarged autolysosomes bearing extremely long, thin LAMP1-positive tubules[3]. Importantly, we also found that *dyn-1(ts)* mutants displayed dramatic tubulation of LMP-1-positive autolysosomes in the *C. elegans* touch neurons (Figure 3B). Taken together our results indicate phylogenetic conservation of function in autolysosome tubule severing mechanisms from nematode to human and suggest that RME-8 is important for autolysosome tubule release.

**Figure 3.**
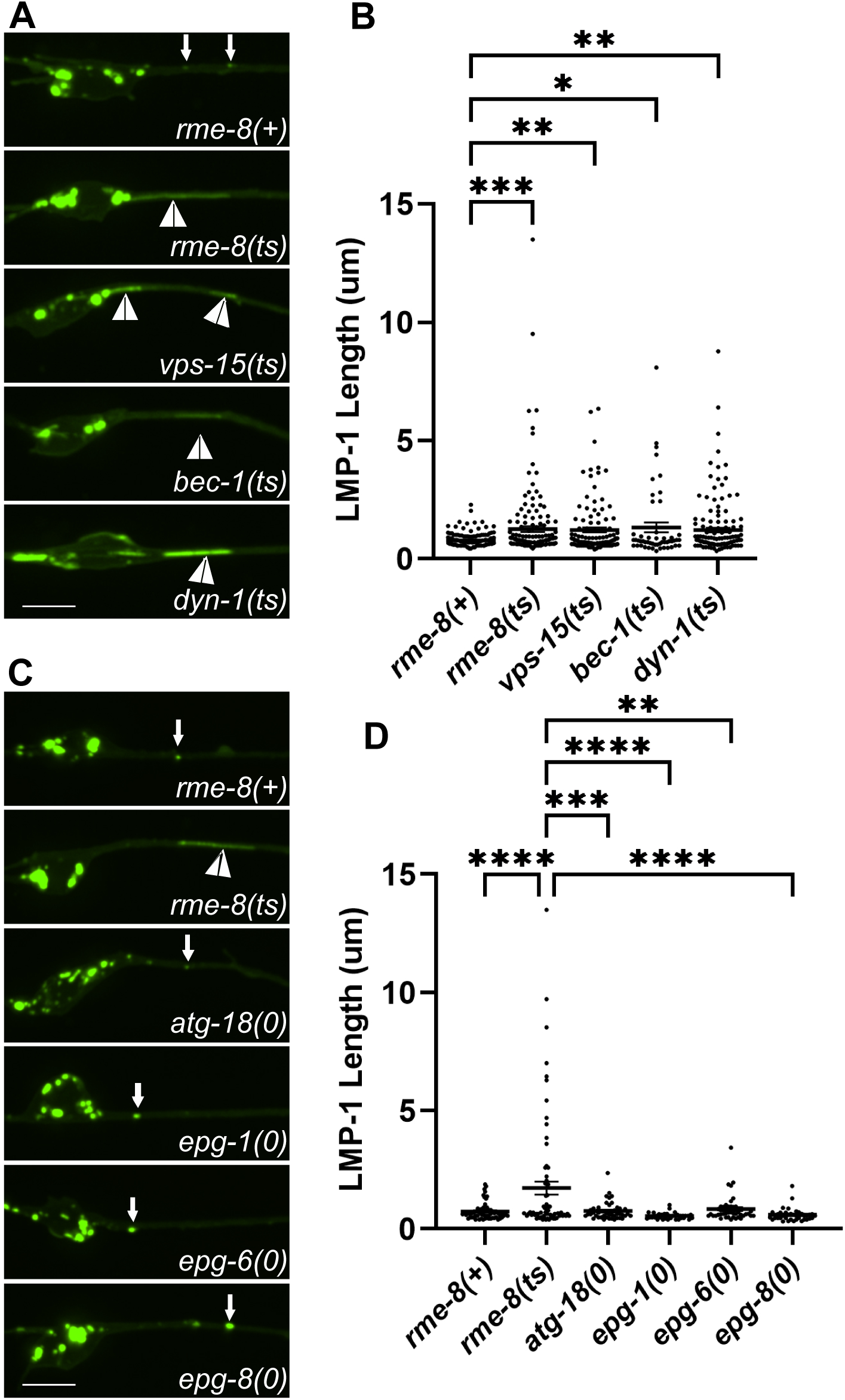
Loss of RME-8 phenocopies autolysosome reformation mutants. **A)** Micrographs are shown of pmec-7 LMP-1::mNG in ALM neurons in *rme-8(+), rme-8(ts), vps-15(ts), bec-1(0),* and *dyn-1(ts)* backgrounds. **B)** Quantification of LMP-1 tubule length (um) in ALM proximal axons in the backgrounds shown in 3A. Each data point represents an individual puncta/tubule. ***P < 0.001, **P < 0.01, *P < 0.05, ns > 0.05 by One-way ANOVA followed with Tukey’s Multiple Comparisons Test. **C)** Micrographs are shown of pmec-7 LMP-1::mNG in ALM neurons in *rme-8(+), rme-8(ts), atg-18(0), epg*-*1(0), epg-6(0), epg-8(0),* backgrounds. Example images and graphs are separate from 3A and 3B because data was obtained during separate experiments. D) Quantification of LMP-1 tubule length (um) in ALM proximal axons in the backgrounds shown in 3C. Each data point represents an individual puncta/tubule. ***P < 0.001, **P < 0.01, *P < 0.05, ns > 0.05 by One-way ANOVA followed with Tukey’s Multiple Comparisons Test.

### Mammalian RME-8/DNAJC13 is also required for neuronal ALR

We also sought to determine if RME-8 (also called DNAJC13) is required for ALR in mammalian neurons. To test this, we analyzed primary cortical neurons derived from the embryonic mouse brain, visualizing autolysosomes with GFP-LAMP1 and mRFP-LC3, knocking down mouse RME­8/DNAJC13 using shRNAs. Importantly, in neurons expressing RME-8/DNAJC13 shRNA, we observed strikingly elongated LAMP1-labeled tubules emanating from vesicles positive for mRFP-LC3, identifying them as autolysosomes (Figure 4A). Under these conditions LAMP1- positive tubule length increased from a mean of <4 um in control shRNA neurons, to ∼18 um in RME-8/DNAJC13 shRNA neurons (Figure 4B). Elongated LAMP1-positive tubules in RME­8/DNAJC13 knock down conditions can be seen in Video S3, and compared to the dynamics of lysosomes in neurons expressing control shRNA (Video S4). The proportion of long tubules was much greater upon knock down of RME-8/DNAJC13, with 39% of tubules having a length of greater than 20um upon loss of RME-8, while 0% of tubules in control cells were longer than 20um (Figure 4C). We also noted that the autolysosomes in RME-8/DNAJC13 knockdown neurons were greatly enlarged, in many cases displaying areas >3 times greater than in controls (Figure 4D), and were fewer in number (Figure4E). This is similar to previously described ALR defects upon loss of known ALR regulators, that also develop enlarged LAMP1-positive autolysosomes due to an inability to recycle material out of the lysosomes after fusion with autophagosomes [1–4]. We also observed a significantly reduced number of LAMP1-positive lysosomes that lack LC3 in RME-8/DNAJC13 knockdown neurons (Figure 4F), further indicating a defect in lysosome reformation from autolysosomes. Taken together our results indicate a clear requirement for RME-8 in completion of ALR in *C. elegans* and mammalian neurons.

**Figure 4.**
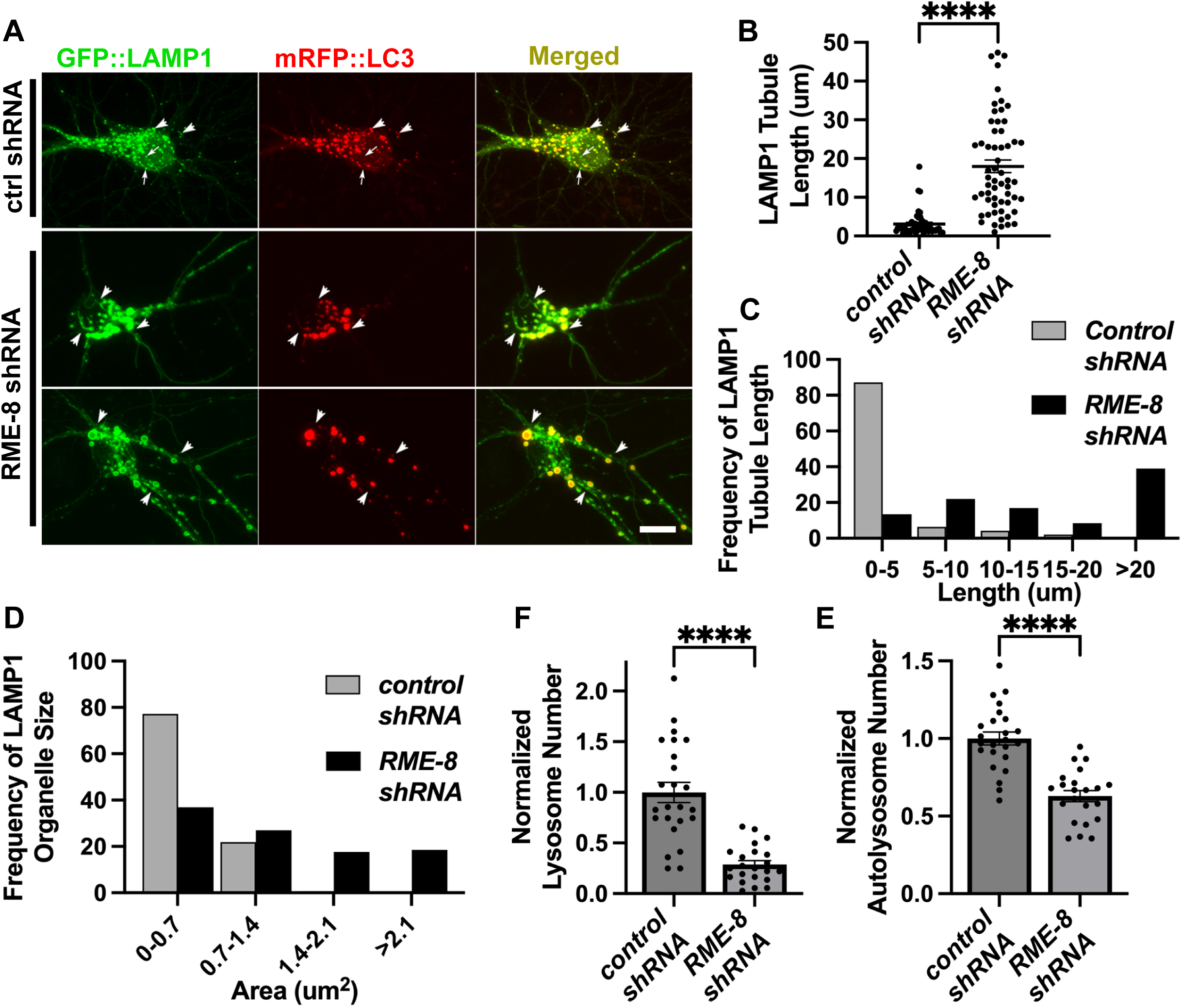
RME-8 knockdown in mouse cortical neurons causes elongated autolysosome tubules and enlarged autolysosomes. A) Micrographs of double-labeled GFP-LAMP1, mRFP-LC3 in primary mouse cortical neurons transfected with control (top panel) or RME8/DNAJC13 (bottom two panels) shRNA. Images are shown for single channel labeling and merged. Arrows (top panel, control shRNA) indicate lysosomes (GFP-LAMP1 positive; mRFP-LC3 negative), arrowheads indicate autolysosomes (GFP-LAMP1 positive; mRFP-LC3 positive). Scale bar, 10 um. B) The average LAMP1 tubules length of each soma is quantified in control and RME­8/DNAJC13 shRNA conditions. Each puncta represents the average length in one neuron. **C)** Frequency distribution of average tubule length per soma in control and RME-8/DNAJC13 shRNA. **D)** Frequency distribution of LAMP-1 positive (lysosomes and autolysosomes) vesicle area per soma in control and RME-8/DNAJC13 shRNA. **B), F), E):** Student’s unpaired t-test. ****P<0.0001.

### RME-8 localization in neurons

RME-8 in neurons could function in ALR directly via a role on autolysosomes, or it could affect ALR indirectly, for instance through endosomes which also fuse with lysosomes, potentially providing regulatory molecules in addition to degradative cargo. To assay RME-8 subcellular localization in *C. elegans* we expressed a rescuing touch-neuron-specific RME-8::GFP transgene. At steady state we did not observe a high degree of RME-8::GFP on LMP-1::mSc-positive lysosomes in the *C. elegans* touch neurons, but some overlap was detected (Figure 5A-B). We observed a high degree of colocalization of RME-8::GFP and endosome marker mSc::SNX-1 in the neuronal soma, similar to our previous results in other cell types (Figure 5A-B). In summary, if RME-8 functions directly on autolysosomes during ALR, it is likely to be transiently associated with a small region of the autolysosome, as has been proposed for proteins like dynamin that participate in autolysosome tubule fission[3].

**Figure 5.**
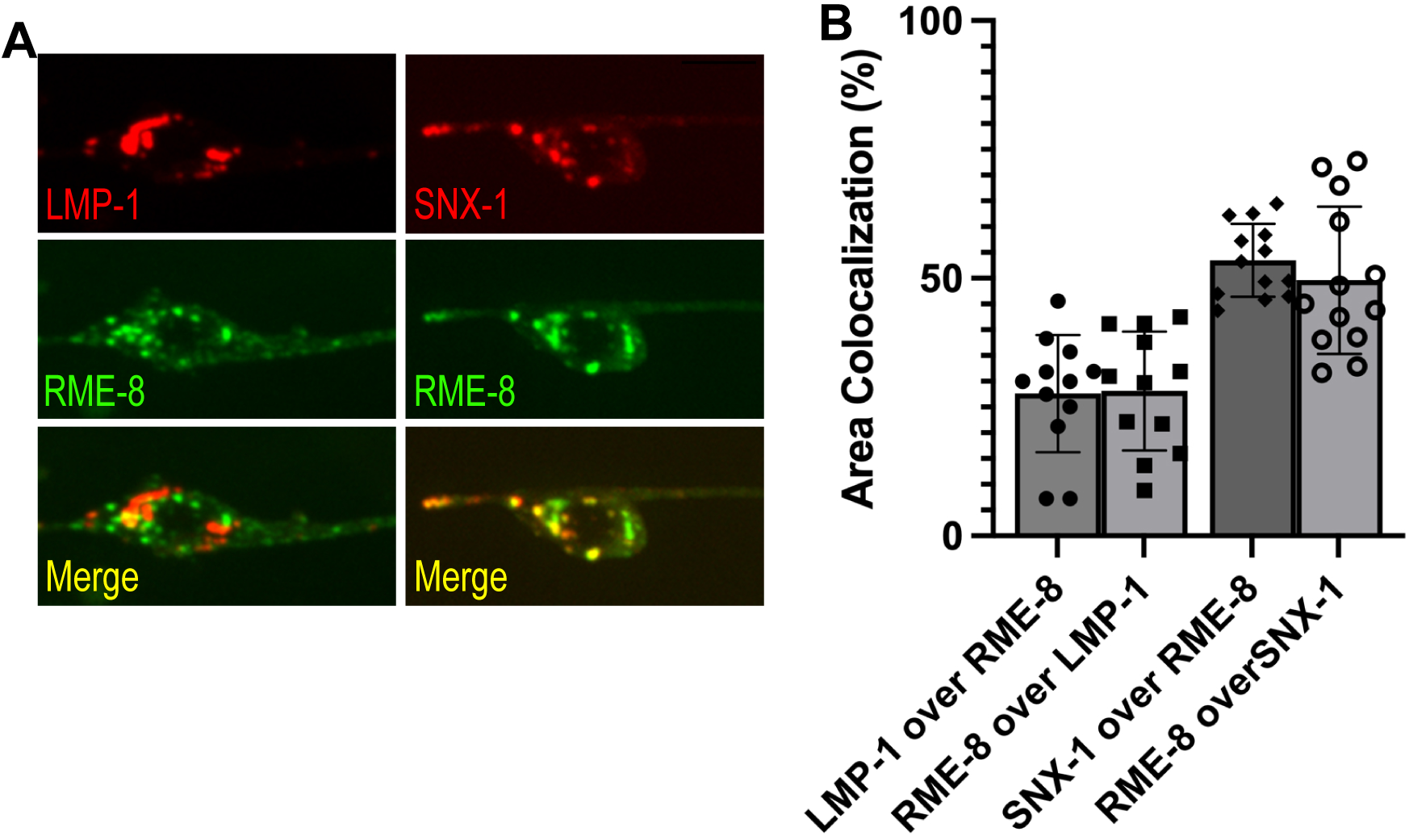
RME-8 localizes primarily to endosomes. **A)** Fluorescent single channel and merged micrographs of double labeled strains LMP-1::mScarlet; RME-8::GFP and mScarlet::SNX-1; RME-8::GFP in wild-type background. Scale bar, 5um. **B)** Quantification of colocalization between strains shown in **A)** using % Area Colocalization.

### Reduced neuronal a utophagv in *C. elegans rme-8* mutants

Given the apparent reduced intensity of mNG::LGG-1 we observed in *rme-8* mutants, we asked if known ALR-relevant proteins are important for maintaining autophagic flux in *C. elegans* neurons. We first investigated this by comparing mNG::LGG-1 levels in touch neuron somata of *rme-8, bec-1, vps-15,* and *dyn-1* mutants (Figure 6A). Beyond ALR, VPS-15 and BEC-1 also impinge upon P1(3)P generation at the omegasome, a pre-autophagosomal structure, but dynamin has not been linked to any direct role in autophagosome formation [44, 45]. We observed a similar decrease in both the integrated and average intensity of mNG::LGG-1 puncta in *rme-8, dyn-1/dynamin, bec-1/beclin,* and *vps-15* mutants (Figure 6B-C). These results are consistent with recent studies linking impaired ALR to decreased autophagy initiation[38].

**Figure 6.**
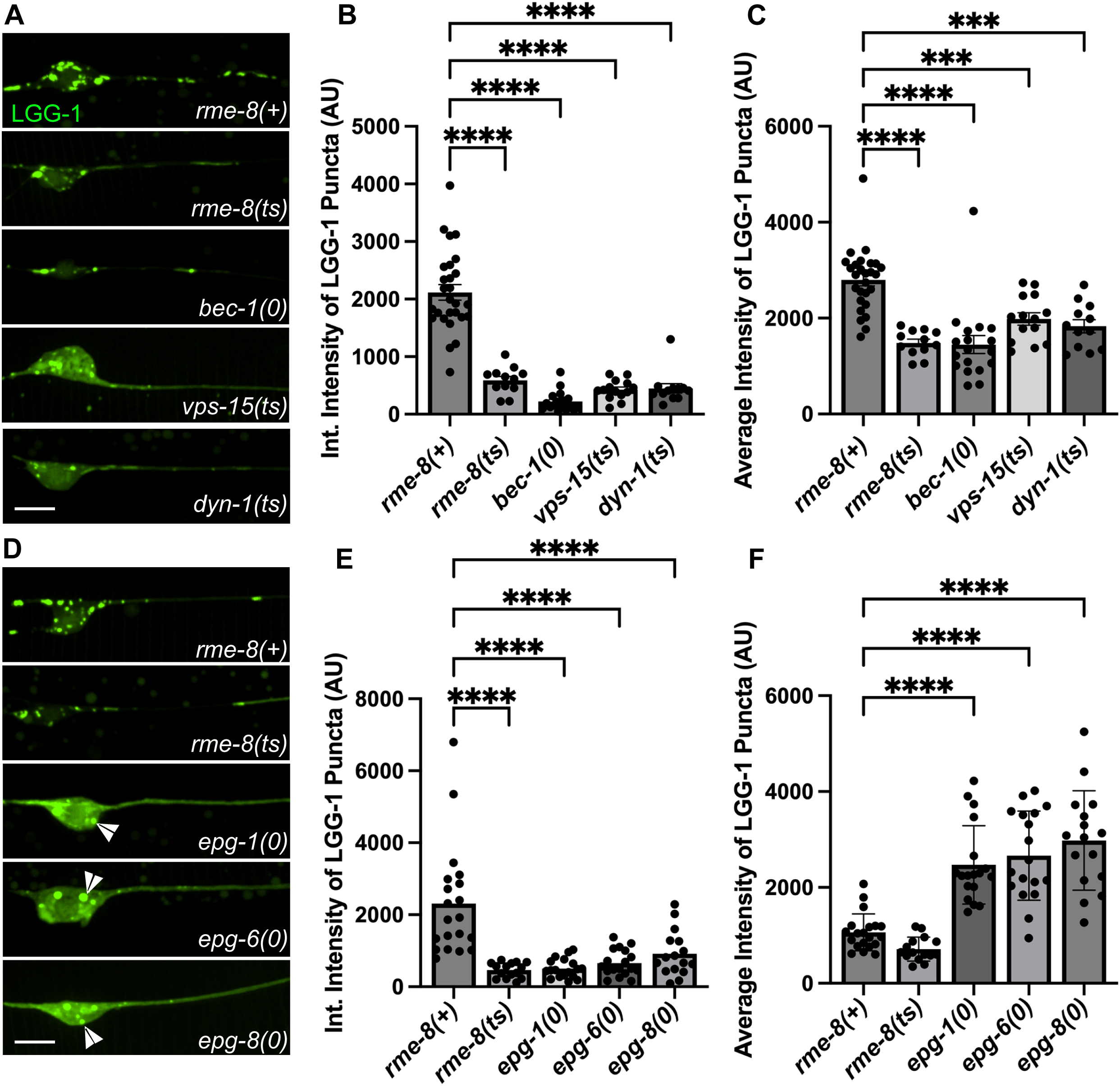
Loss of RME-8 causes decreased autophagosome levels like known ALR mutants. **A), D)** Micrographs of pmec-7 LGG-1::mNG in C. *elegans* ALM neurons. Area shown includes soma (left) and proximal axon (right). Example images and graphs are separate because data was obtained during separate experiments. Scale bars, 5 um. **B), E)** Quantification of integrated intensity of LGG-1::mNG puncta in backgrounds shown in **A)** and **D)** (AU; arbitrary units). **C), F)** Average Intensity of LGG-1 positive puncta in axons and somas of backgrounds shown in **A)** and **D). ***P <** 0.001, ****P < 0.0001 by One-way ANOVA followed with Tukey’s Multiple Comparisons Test.

We also examined mNG::LGG-1 levels and distribution in *epg-1/atg13-like*[46]*, epg-6/atg18- like*[47], and *epg-8* mutants[48]. These proteins have clearly defined roles within the canonical autophagic initiation pathway, but without known effects on ALR. We found that *epg-1, epg-6,* and *epg-8* mutants displayed a strongly diffusive mNG::LGG-1 signal in the touch neuron soma, with two or three apparent large aggregates near the nucleus (Figure 6D; white arrowheads). While *epg-1, epg-6,* and *epg-8* all have decreased integrated intensities (Figure 6E), similar to ALR mutants, the average intensity of the puncta is significantly higher than wild-type puncta (Figure 6F), in stark contrast to the decreased average intensity of ALR mutants (Figure 6C). This aggregation phenotype has been previously observed in cells expressing tagged LGG-1 in which autophagy initiation is strongly blocked, as expected in these mutants, resulting in overexpressed LGG-1 accumulating in an unprocessed form in aggregates[46, 49, 50]. This is quite different from the reduced levels of mNG::LGG-1 fluorescence we observed in *rme-8, dyn­1, bec-1,* and *vps-15* mutants, where dimmer mNG::LGG-1 is found in more numerous small puncta within the touch neuron soma (Figure 6A). Taken together we interpret the *rme-8* mutant phenotype as reduced but not blocked neuronal autophagy, potentially as an indirect consequence of defective ALR.

Another possibility is that defective autophagy initiation is sufficient to produce a rme-8-like lysosome phenotype with accumulation of long lysosomal tubules. However, when we measured LMP-1::mNG tubule length in *epg-1(0), epg-6(0), epg-8(0)* and *atg-18(0)* mutants in the touch neurons, we did not find any significant elongation of LMP-1 tubules (Figure 3C-D). These results suggest that a simple block in autophagy initiation is not sufficient to produce this phenotype.

### Neuronal autophagic flux requires RME-8/DNAJC13 in mammalian neurons

Given our results in *C. elegans,* we examined the effects of RME-8/DNJC13 on autophagic flux in primary mouse cortical neurons. Similar to our observations in *C. elegans* neurons, we found that autophagic vacuole (AV) density per soma, as visualized by GFP-LC3, was decreased in RME-8/DNAJC13 shRNA neurons compared to controls (Figure 7A, quantified in 7B). We also measured AV density in the presence of pepstatin and E64D, lysosomal protease inhibitors, which block the autolysosome-mediated degradation of autophagic cargo, including GFP-LC3. This type of analysis is often used to judge levels of autophagic flux, since reduced GFP-LC3 levels can result from reduced autophagy, or increased lysosomal degradation of autophagic cargo. If autophagic flux is reduced, then GFP-LC3 levels should also be reduced, even when lysosomal degradation is blocked. However, if GFP-LC3 levels are reduced due to increased lysosomal activity, blocking such lysosomal activity should block the observed GFP-LC3 reduction. We found GFP-LC3 AV density was still decreased in RME-8 shRNA neurons treated with pepstatin (Fig 7A-B). We interpret these results to indicate reduced autophagic flux in RME-8/DNAJC13-deficient mammalian neurons, potentially at an early stage in autophagy initiation or autophagosome formation.

**Figure 7.**
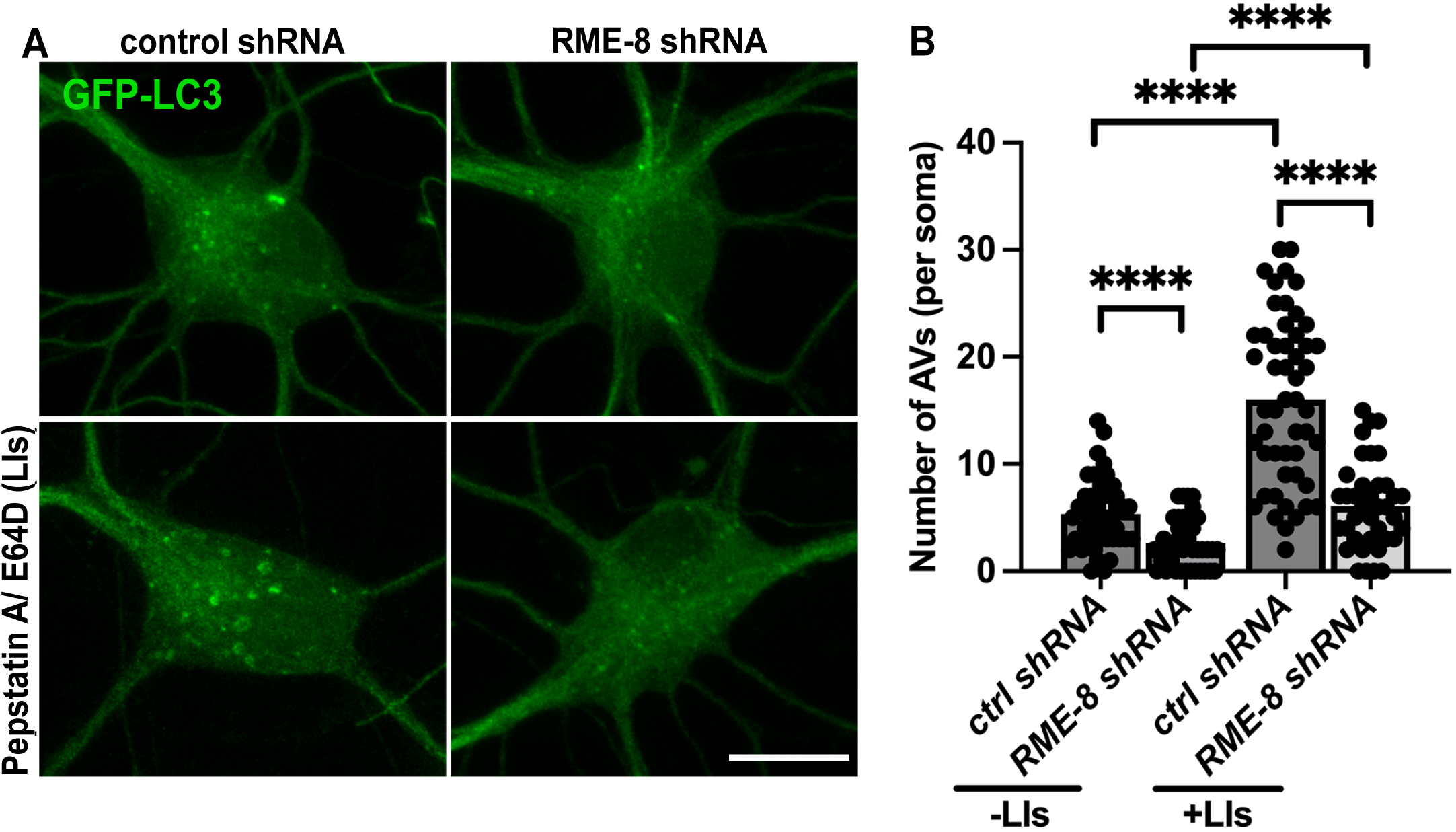
RME-8/DNAJC13 knock down in mouse cortical neurons reduces autophagic flux. **A)** Micrographs are shown of *in vitro* mouse cortical neurons transfected with GFP-LC3. A 2x2 factorial design was used, conditions were treatment with either control or RME-8/DNAJC13 shRNA, and +/- pepstatin/ E64D (lysosome inhibitors; LIs). Scale bar, 10 um. **B)** Autophagic vacuole (AV) density is quantified in neurons for the 2x2 factorial design shown in **A). ****P <** 0.0001 by Student’s t-test.

### RME-8, BEC-1/beclin, and VPS-15 are required for efficient clathrin recruitment to a uto lvsosomes

One key molecule associated with the biogenesis of autolysosome reformation tubules is clathrin[2, 4, 38]. Previously, we and other groups had identified an important role for RME-8 in controlling clathrin levels on endosomes in non-neuronal cells [23, 51]. Thus, we hypothesized that RME-8 activity might be important for controlling lysosomal clathrin levels relevant to ALR in neurons. To test this idea we established touch neuron-specific lines expressing clathrin light chain tagged with mNeonGreen (CLIC-1::mNG). We found that CLIC-1::mNG was clearly enriched on LMP-1::mSc labeled lysosomes, supporting a conserved role for clathrin in lysosome homeostasis (Figure 8A-B). Furthermore, we found that *bec-1* and *vps-15,* mutants that accumulate long autolysosomal tubules like *rme-8,* displayed significantly less recruitment of CLIC-1::mNG to neuronal LMP-1-positive lysosomes than controls. Importantly, we found that *rme-8* mutants display an even more severe deficit in CLIC-1::mNG recruitment to LMP-1- positive lysosomes than *bec-1* or *vps-15* mutants (Figure 8A-B). Notably, the severely elongated LMP-1::mSc tubules in *rme-8(b1023ts)* mutants completely lack CLIC-1::mNG labeling. We also identified a defect in clathrin recruitment to neuronal lysosomes in *snx-1* mutants (Figure 8A-B). These mutants are missing RME-8 partner protein SNX-1 (Sorting Nexin 1) that promotes RME-8 activity [15, 24]. BEC-1 and VPS-15 may also be required for efficient RME-8 function, since RME-8 has been reported to have an N-terminal P1(3)P binding domain[21, 51]. Overall, we find that RME-8, SNX-1, BEC-1, and VPS-15 are required for efficient recruitment of clathrin to neuronal lysosomes. Since clathrin has been reported to be a key molecule in ALR tubule biogenesis, reduced levels of clathrin on lysosomes may lead to defective ALR.

**Figure 8.**
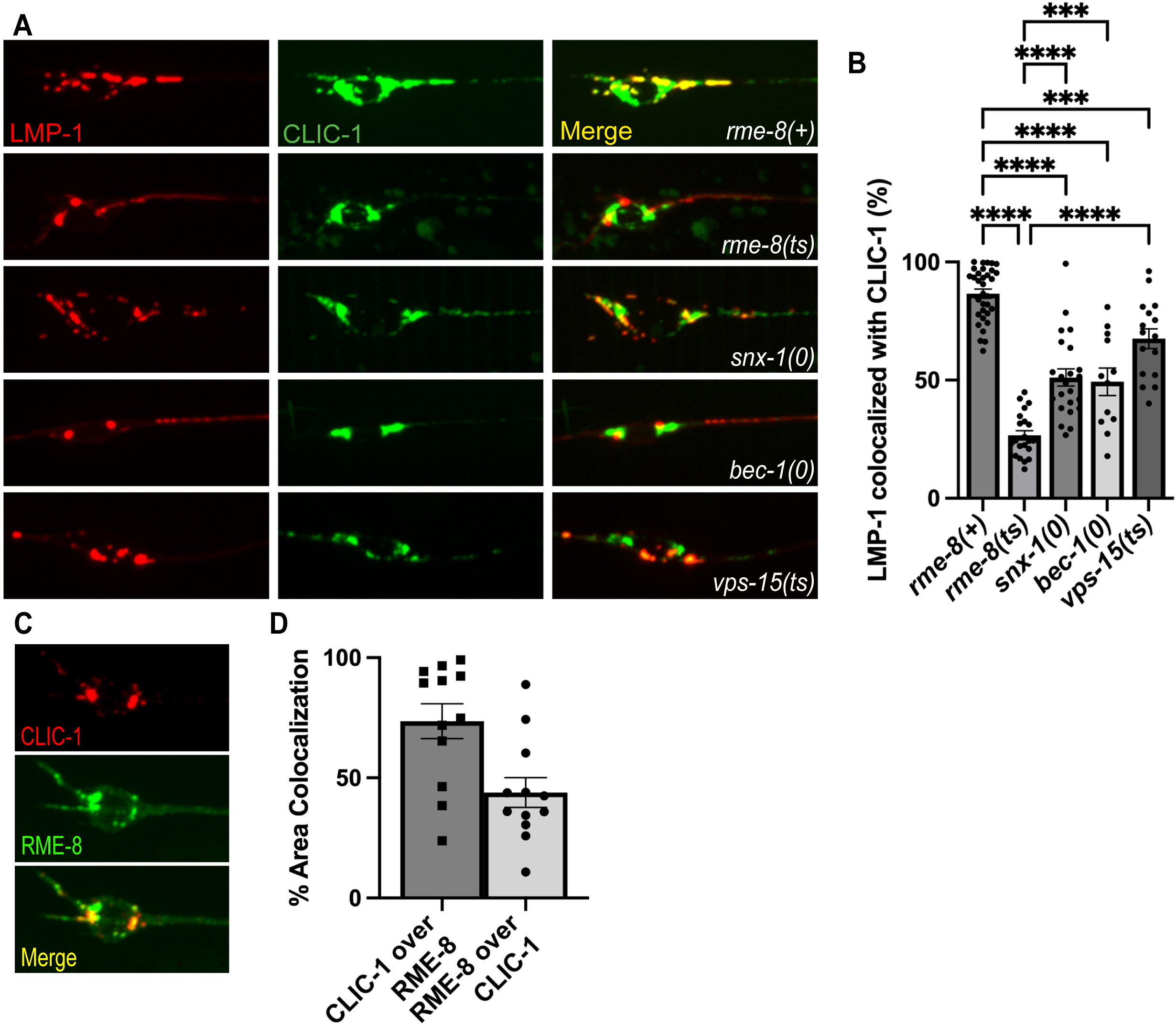
RME-8 is required for efficient clathrin recruitment to lysosomes. **A)** Single-channel and merge images of double-labeled LMP-1::mSc and CLIC-1::mNG in C. elegans’ ALM neurons. **B)** Quantification of the percent of LMP-1::mSc that has CLIC-1::mNG present on it. Scale bar, 5 um. One-way ANOVA followed with Tukey’s Multiple Comparisons Test. *P < 0.05, **P < 0.01, ***P < 0.001, ****P < 0.0001.

## DISCUSSION

Autophagic engulfment of cytoplasmic content represents a major pathway required for maintaining cellular proteostasis and organelle integrity, as well as survival under nutrient deprivation conditions. Autophagy is likely to be especially important in neurons, which tend to be extremely long-lived and cannot divide, and thus lack an effective mechanism to dilute protein aggregates and defective organelles as rapidly dividing cells can. Consistent with this idea, neurons maintain a relatively high basal rate of autophagy, and autophagic dysfunction has been repeatedly implicated in neurodegenerative disease (reviewed in [7, 9, 37, 52, 53]).

Once autophagosomes engulf their cargo they must fuse with lysosomes, producing autolysosomes that degrade their contents. Maintaining a pool of active lysosomes to receive autophagic cargo requires efficient recovery of membrane and other machinery from autolysosomes to reform fresh lysosomes. Defective autolysosome reformation is emerging as an important disease mechanism in its own right, associated with inflammatory and liver disease, muscular dystrophy, metabolic syndrome, Charcot-Marie-Tooth disease, lysosome storage disease, and neurodegenerative disorders such as hereditary spastic paraplegia and Parkinson’s disease.

Given the genetic connections suggesting linkage of human RME-8/DNAJC13 to Parkinson’s disease and essential tremor, we sought to analyze the requirement for RME-8 in *C. elegans* and mouse neurons. Our experiments identified an important and conserved role of RME-8 in autolysosome reformation in both systems. Loss of RME-8 produced enlarged autolysosomes bearing persistently elongated membrane tubules indicative of a defect in ALR tubule fission to release protolysosomes. In the case of RME-8/DNAJC13 depleted mouse cortical neurons we could document a loss of LC3-negative lysosomes, further indicating a failure in lysosome reformation. We also noted in *C. elegans* bearing an endogenous *rme-8(N861S)* allele, equivalent to human PD-associated allele N855S, an age-dependent effect on RAB-7-positive tubule accumulation, suggested that disease effects could derive from weak dysfunction in ALR.

Extended analysis in the *C. elegans* system revealed a striking loss of clathrin from neuronal autolysosomes, suggesting that clathrin malfunction may underlie the ALR defects when RME-8 is missing. Interestingly we also observed reduced lysosome-associated clathrin in mutants lacking RME-8 partner SNX-1, as well as mutants lacking P13-kinase complex proteins VPS-15/P150 and BEC-1/beclin. While P1(3)P generated by Vps34 under the control of VPS-15 and BEC-1 homologs was previously linked to ALR, an effect on clathrin was unexpected, since autolysosomal clathrin is thought to assemble with the AP2 clathrin adapter on P1(4,5)P2 enriched membranes. However, RME-8 is known to bind to P1(3)P and to affect clathrin dynamics, likely due to its activity as a Hsc70 co-chaperone, so RME-8 could be a key effector of P1(3)P during ALR[23, 24]. Previous work indicated that *bec-1* mutants affected RME-8 recruitment to membranes in the *C. elegans* intestine [42]. A complex containing ZFYVE26, SPG11, and AP-5 has been proposed as a P1(3)P effector during ALR, but *C. elegans* lacks homologs for any of these proteins, so they cannot explain the conserved requirement from *C. elegans* to mammals for P1(3)P during ALR[11, 54].

Elegant experiments by Rong et al., 2012 suggested dual roles for clathrin in ALR[2]. These authors found that when they depleted P1(4,5)P2 and clathrin from the vesicular body of the autolysosome, tubule budding failed. Alternatively, when P1(4,5)P2 and clathrin were depleted from the tubule, tubule fission failed, blocking protolysosome release. Other work also implicates clathrin regulation in ALR. For instance, abnormally high autolysosome P1(4,5)P2 levels were found to result in increased clathrin on elongated ALR tubules, suggesting that general dysregulation of clathrin on tubules may be sufficient inhibit tubule release.

While we did not observe high-level constitutive residence of RME-8 protein on lysosomes, RME-8 could be present quite transiently, as are some proteins that act in late stages of vesicle budding. If so, RME-8 could play a direct role in clathrin dynamics regulating ALR. While DNAJ­domain co-chaperone auxilin is best known for uncoating clathrin from clathrin-coated vesicles, *in vitro* auxilin promotes clathrin assembly, and has also been proposed to provide co-chaperone activity that helps rearrange clathrin coats from flat to curved during budding[55]. RME-8 could act similarly on autolysosomes. RME-8 could also play an indirect role via global clathrin metabolism, as our previous work identified a requirement for RME-8 in uncoating clathrin reservoirs on endosomes, such that loss of RME-8 activity could reduce soluble, cytosolic clathrin levels in the cell needed for assembly on autolysosomes. More analysis will be required to distinguish between these possibilities.

While it is not clear how clathrin contributes to tubule release, clathrin/AP2 assembly likely acts as a precursor to the recruitment of dynamin, the pinchase thought to directly sever ALR tubules, as it does during clathrin-mediated endocytosis[56]. Effects on dynamin may also underlie some of the effects on ALR that we observed in mutants defective in P13-kinase regulators BEC-1/beclin and VPS-15/P150. Munson et al found that when a lysosomal pool of P1(3)P is reduced by inhibition of the Vps34 complex, long ALR tubules accumulate[6, 57]. P1(3,5)P2 produced from P1(3)P on late endosomes by the PIKFyve complex is also important for neuronal lysosome function and may contribute to ALR [58, 59]. It is unclear to what extent dynamin requires P1(3)P and/or P1(3,5)P2 for its function, but *in vitro* the dynamin PH-domain binds multiple phosphoinositides, including P1(3)P, P1(3,5)P2, and P1(4,5)P2 [60]. Furthermore, *in vitro* P1(3)P and P1(3,5)P2 shows high efficiency in stimulating assembly-dependent GTPase activity by dynamin in complex with its partner Snx9, which also displays broad phosphoinositide binding [61]. Phagocytosis studies have expanded upon this idea *in vivo,* indicating a need for precise and sequential regulation of P1(4,5)P2 and P1(3)P in phagosome release from the plasma membrane mediated by dynamin and Snx9/LST-4[62–64]. Similar regulation may be required on ALR tubules for dynamin-mediated protolysosome release.

In addition to effects on ALR, we also noted an apparent defect in autophagy, with strongly reduced levels of LGG-1/LC3 in the neuronal soma of *rme-8, bec-1/beclin, vps-15/P150,* and *dyn-1/dynamin* mutants. This was particularly surprising in the case of DYN-1, since dynamin has not been proposed to have any role in autophagosome formation. Our data argue against the ALR tubule accumulation defects we observed for these mutants being an indirect effect of reduced autophagic flux, as we did not observe tubule accumulation in mutants that affect autophagy initiation but that have no known role in ALR. Results vary in the literature as to whether ALR is required to maintain autophagosome formation rates, but our data supports such a requirement. It seems likely that ALR is required to recycle some component(s) or regulator(s) of autophagy from autolysosomes, and that extended failure in ALR leads to reduced rates of autophagy. An interesting candidate for a recycling cargo during ALR is Atg9, the only transmembrane regulator of autophagy initiation. Atg9 was recently shown to be recycled from autolysosomes by the recycler complex, and RME-8 has also been implicated in Atg9 trafficking [65, 66]. It will be of great interest to further analyze the mechanism of RME-8 function in ALR, and to gain further insight into mechanisms by which ALR could influence autophagic flux.

## MATERIALS AND METHODS

All *C. elegans* strains were derived originally from the wild-type Bristol strain N2. Worm cultures, genetic crosses, and other *C. elegans* husbandry were performed according to standard methods[67]. Since the experiments utilized temperature sensitive strains, all strains were maintained at 15°C. A complete list of strains used in this study can be found in Table S1. Every strain listed was created in the Grant Lab for use in this study.

### Construction of *C. elegans* plasmids

*C. elegans* expression plasmids utilized the Pmec-7 promoter from the *mec-7* gene for touch neuron expression[68]. Vector details are available upon request. Cloning was performed using the Gateway *in vitro* recombination system (Invitrogen, Carlsbad, CA) using Grant lab-modified versions of MiniMos enabled vectors pCFJ1662 (Hygromycin resistant; Addgene #51482) and pCFJ910 (G418 resistant; Addgene #44481) (gifts of Erik Jorgensen, University of Utah): pCFJ1662 Pmec-7 GTWY mNeonGreen 1et858 (34F6); pCFJ1662 Pmec-7 mNeonGreen GTWY 1et858 (34D4); pCFJ1662 Pmec-7 GTWY oxGFP 1et858 (36G3); pCFJ910 Pmec-7 mScarleti GTWY 1et858 (33B6); and pCFJ910 pmec-7 GTWY mScarleti (35D2). pDONR221 entry vectors containing coding regions for *rab-7, Imp-1, lgg-1, clic-1, snx-1, rme-8,* were recombined into neuronal destination vectors by Gateway LR clonase II reaction to generate C-/N- terminal fusions. Single-copy integrations were obtained by MiniMOS technology [69].

### Image acquisition and Analysis in *C. elegans*

Prior to imaging, L4 hermaphodites maintained at 15°C were picked to fresh OP50 plates and plates were temperature shifted to 25°C (nonpermissive temperature for *rme-01023ts1)* for 24-30 h prior to imaging. Live animals were picked into 2 uL of 10 mM levamisole on the center of a cover slip. After two minutes, 5% agarose pads were placed on top. Fluorescence images were obtained using a spinning-disk confocal imaging system: Zeiss Axiovert Z1 microscope equipped with X-Light V2 Spinning Disk Confocal Unit (CrestOptics), 7-line LDI Laser Launch (89 North), Prime 95B Scientific CMOS camera (Photometrics) and oil-immersion objective (100X). Fluorescence images were captured using Metamorph 7.10 software. Z series of 25-30 optical sections were acquired using a 0.2[1m step size in order to image through the entire soma. Exposure times ranged from 50ms to 300 ms based on the fluorescent protein being imaged such that images acquired were at minimum in the 12-bit and did not become saturated in the 16-bit range in any genetic background.

### Intensity and Length measurements

All data quantification was done using Metamorph 7.10 and data was recorded into Excel. Maximum projections were generated from Z series captured under identical acquisition parameters for each experiment. For each experiment, maximum projections were scaled identically prior to thresholding or length analysis. Puncta within the soma, dendrite, and proximal axon (within ^—^30um of the soma) were manually thresholded and the average, maximum, and integrated intensity measurements recorded using Metamorph’s “Region Measurements” tool. Each worm was treated as one data point.

Maximum projections were also used for length measurements. Lengths were measured and recorded using Metamorph’s “region measurements” and using the line tool to trace the length of every puncta/ tubule in the axon. The length of every visible puncta/ tubule within each axon was measured and counted as a data point. Analysis of tubule elongation was restricted to axons due to the poor spatial resolution of puncta within the soma. For each experiment and genetic background, 15-25 animals were analyzed.

Colocalization analysis was performed using MetaMorph 7.7 colocalization plugin, whereby intensities in each channel were thresholded and analyzed.

### Mouse cortical neuron plasmids and reagents

DNAJC13 shRNA plasmid(sc-77968-SH) and Control shRNA Plasmid-A(sc-108060) were from Santa Cruz Biotechnology. GFP-LC3B, GFP-LAMP1, and mRFP-LC3 constructs were prepared as previously described[70–76]). E64D (Cat. 330005), and Pepstatin A (Cat.516481) (Sigma).

### Transfection of cultured cortical neurons

Cortices were dissected from E18–19 mouse embryos as described[77–79]). Cortical neurons were dissociated by papain (Worthington, Lakewood, NJ) and plated at a density of 100,000 cells per cm^2^ on polyornithine- and fibronectin-coated coverslips. Neurons were grown overnight in a plating medium (5% FBS, insulin, glutamate, G5, and B27) supplemented with 100× L-glutamine in Neurobasal medium (Invitrogen). Starting at DIV2, cultures were maintained in a conditioned medium with half-feed changes of neuronal feed (B27 in

Neurobasal medium) every 3 days. Neurons were co-transfected with Control shRNA (Cat. sc108060) or DnaJC13 (RME-8) shRNA (Santa Cruz, Cat. sc-77968-SH) with GFP-LC3, GFP-LAMP1, or GFP-LAMP1 and mRFP-LC3 at DIV5 using Lipofectamine 2000 (Invitrogen) before live-cell imaging 7–12 days after transfection before quantification analysis. For some experiments, 24-hour incubation with lysosomal inhibitors E64D (40 μM) and Pepstatin A (40 μM) was applied to suppress lysosomal degradation.

### Image acquisition and quantification in cultured neurons

For live-cell imaging, cells were transferred to Tyrode’s solution containing 10 mM HEPES, 10 mM glucose, 1.2 mM CaCl_2_, 1.2 mM MgCl_2_, 3 mM KCl, and 145 mM NaCl, pH 7.4. The temperature was maintained at 37°C with an air stream incubator. Confocal images were obtained using an Olympus FV3000 oil immersion 60x oil immersion lens (1.3 numerical aperture) with a sequential-acquisition setting, using 488 nm excitation for GFP-LC3 or GFP­LAMP1 and 543 nm for mRFP-LC3. Images were acquired using the same settings below saturation at a resolution of 1,024 x 1,024 pixels (8-bit). Eight to ten sections were taken from the top-to-bottom of the specimen and brightest point projections were made. Time-lapse sequences of 1,024 x 1,024 pixels (8 bit) were collected at 2 s intervals with 1% intensity of the laser to minimize laser-induced bleaching and cell damage while maximizing pinhole opening. Time-lapse images were captured at a total of 100 frames. All recordings started 6 min after the coverslip was placed in the chamber. Images were imported to Adobe Photoshop and morphometric measurements were performed using NIH ImageJ. The thresholds in all images were set to similar levels. Measured data were imported into Excel software for analysis. Data were obtained from at least three independent experiments.

### Graphing and Statistics

All results were graphed using GraphPad Prism 9 software, generating scatter plots depicting individual data points. Bars represent mean with SEM. Significance was measured by Student’s unpaired t-test when comparing 2 strains/conditions or Ordinary one-way ANOVA when comparing >2 strains, unless otherwise noted. Data were considered statistically different at P < 0.05. P < 0.05 is indicated with single asterisks, P < 0.01 with double asterisks, P < 0.001 with triple asterisks, and P < 0.0001 with quadruple asterisks.

## Acknowledgments

We would like to thank the Grant lab for critical comments and suggestions. We thank Peter Schweinsberg and Ge Bai for technical assistance in plasmid construction, and Helen Ushakov for expert microinjection. We thank Alicia Melendez for expert advice and ideas. This work was supported by NIH Grant 5R01GM135326 to B.D.G.

---

**Movie S1. Dynamics of LMP-1::mNG labeled lysosomes in *rme-8(ts) C. elegans* ALM soma shows persistent lysosomal tubulation.** Time lapse image sequences of ALM neurons expressing LMP-1 are shown. An elongated tubule emanating from a lysosome vesicle can be seen. The tubules expands and contracts, but does not sever. Images were collected at 1s increments for 45s.

**Movie S2. Dynamics of LMP-1::mNG labeled lysosomes in wild-type *C. elegans* ALM soma.** Time lapse image sequence of ALM neurons expressing LMP-1 in an *rme*-*8(+)* background is shown. A lysosome in the lower portion of the soma can be observed producing tiny transient tubular processes. Images were collected at 1s increments for 90s.

**Movie S3. Dynamics of lysosomal organelles marked by GFP-IAMP1 in the proximal processes of the cortical neuron expressing DNAJC13(RME-8) shRNA.**

Cortical neurons were transfected with GFP-LAMP1 and RME-8/DNAJC13 shRNA at DIV5, followed by time-lapse imaging at DIV16. Note that GFP-LAMP1-indicated lysosomal organelles appear as elongated tubules and undergo oscillatory movement in proximal processes to the neuronal soma (right). Time-lapse sequences were collected at 2-sec intervals during a 200-s observation.

**Movie S4. Dynamics of GFP-LAMP-1-labeled lysosomes in the proximal processes of control cortical neuron expressing Control shRNA.**

Cortical neurons were transfected with GFP-LAMP1 and Control shRNA at DIV5, followed by time-lapse imaging at DIV16. Note that lysosomal organelles appear as small puncta in the neuronal process and undergo dynamic movement in a predominant retrograde direction toward the soma (left). Time-lapse sequences were collected at 2-sec intervals during a 200-s observation.

**Supplementary Table 1:**
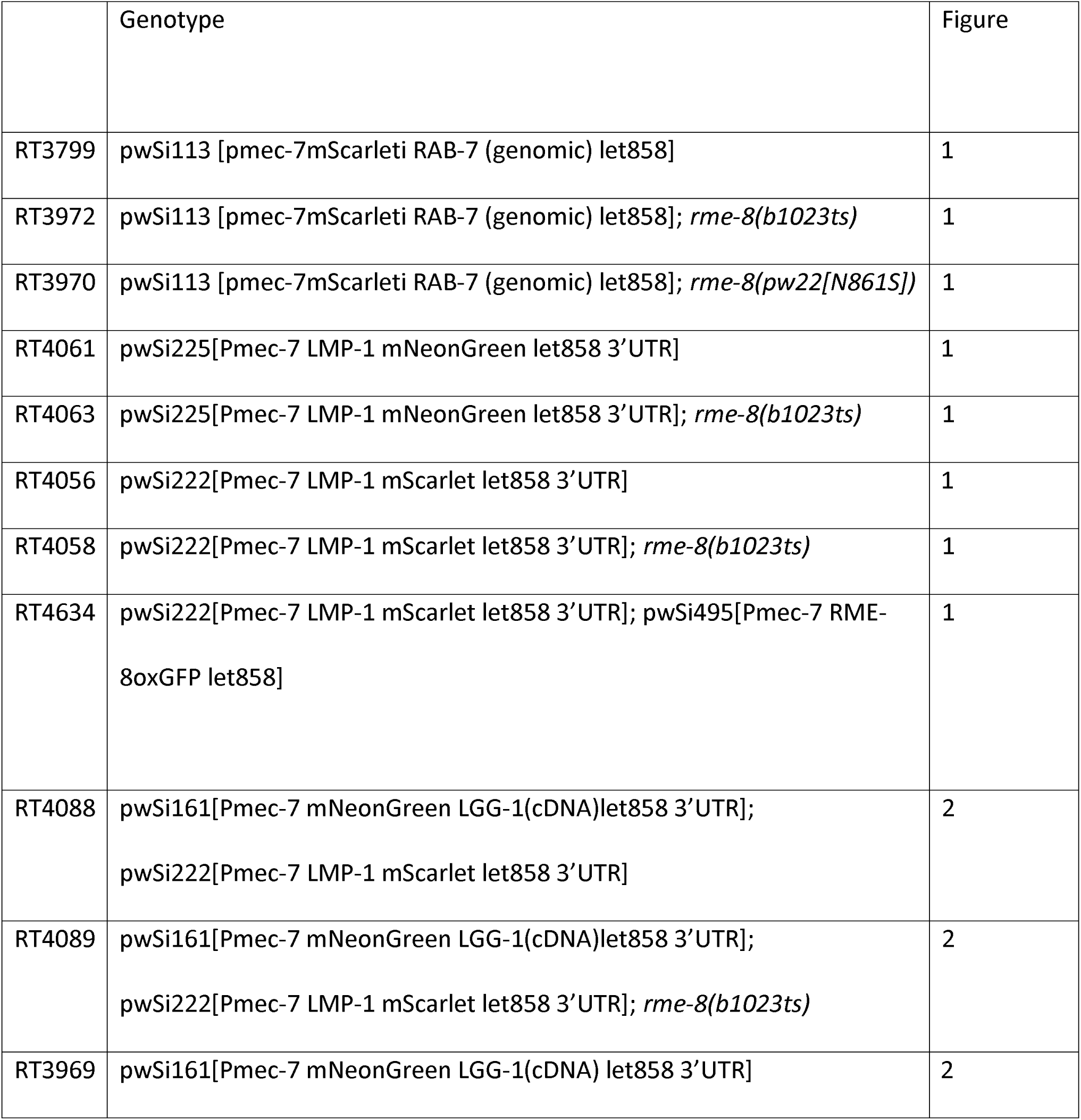
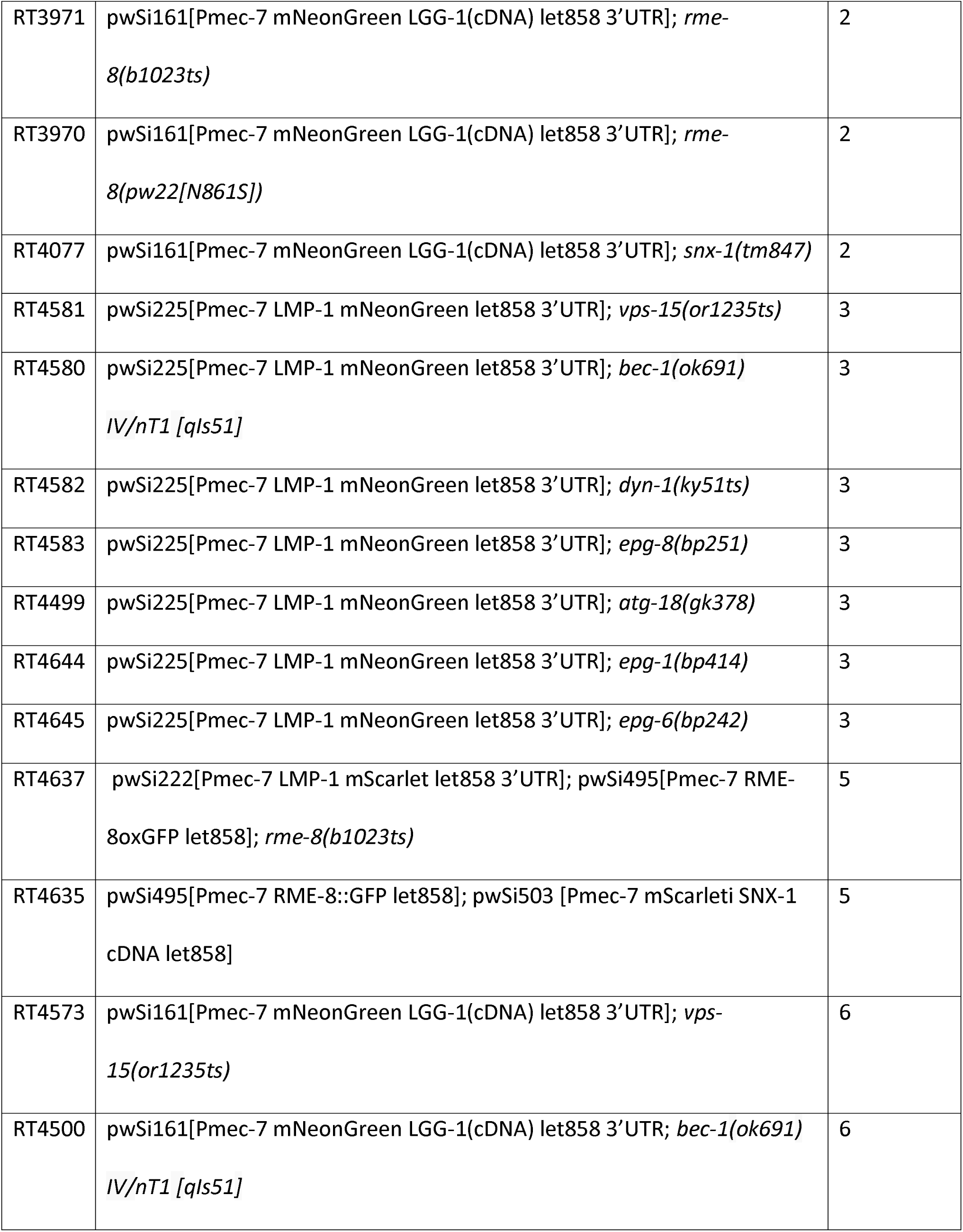
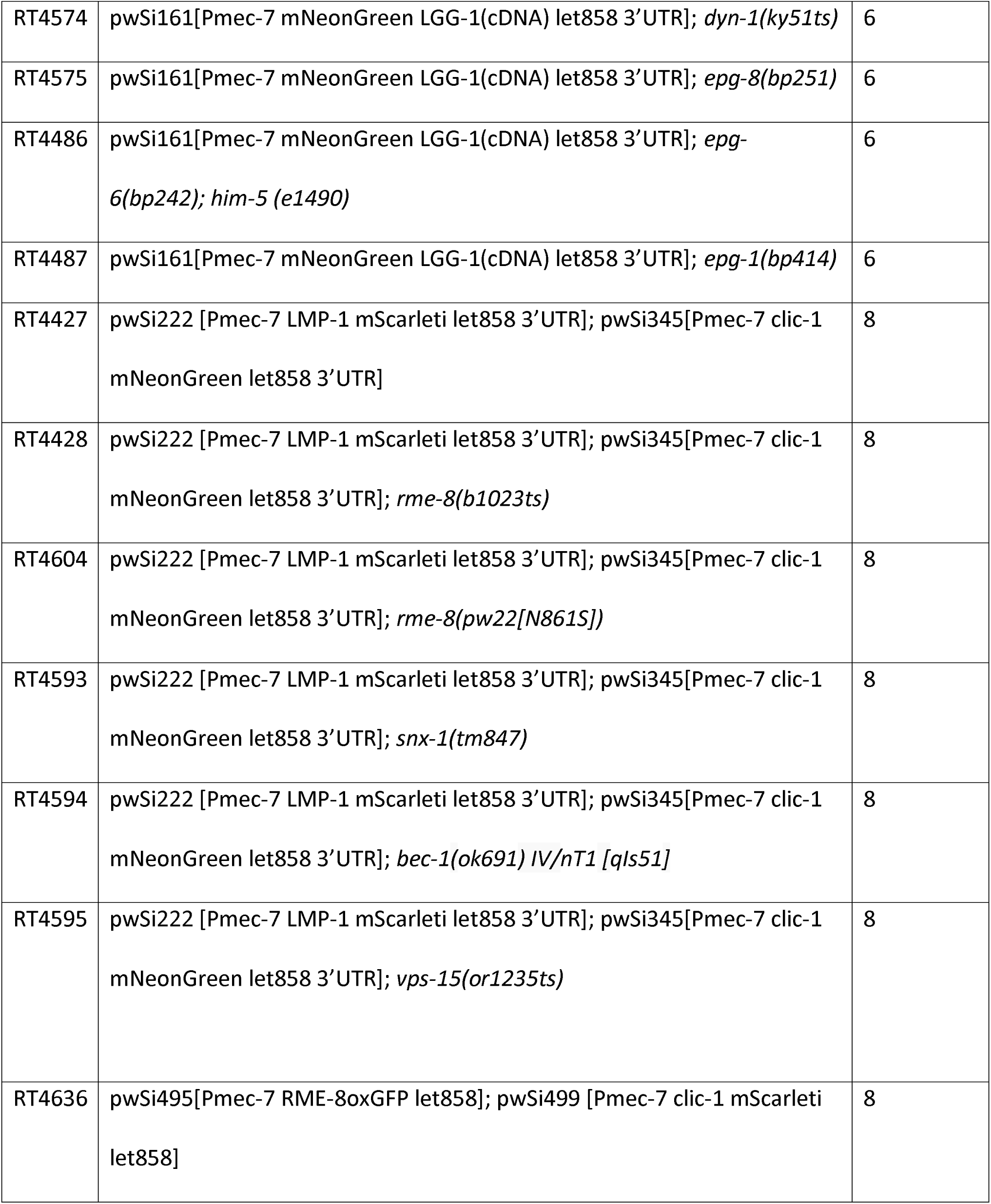
*C. elegans* Strains used in this study

## Notes

### Competing Interest Statement

The authors have declared no competing interest.

